# The Population Structure and Selection Signal Analysis of Shenxian Pigs based on Genome Resequencing Technology

**DOI:** 10.1101/2021.02.08.430358

**Authors:** Liu Diao, Lu Chunlian, Li Shang, Jia Mengyu, Li Sai, Ren Liqin, Miao Yutao, Cao Hongzhan

## Abstract

Shenxian pigs are the only local black pig of Hebei Province, and were listed in the Genetics of Livestock and Poultry Resources of China in 2016. This breed of pig is considered to be a valuable local pig germplasm genetic resource in China. In the present study, in order to understand the genetic variations of Shenxian pigs, identify selected regions related to superior traits, and accelerate the breeding processes of Shenxian pigs, the whole genome of the Shenxian pigs was resequenced and compared with that of large white pigs. The goal was to explore the germplasm characteristics of Shenxian pigs.The results obtained in this research investigation revealed that the genetic relationships of the Shenxian pig breed were complex, and that sub-populations could be identified within the general population. A total of 23M SNP sites were obtained by whole genome resequencing, and 1,509 selected sites were obtained via bioinformatics analyses. It was determined after annotation that a total of 19 genes were enriched in three items of bioengineering, molecular function, and cell composition.During this research investigation, the aforementioned 19 genes were subjected to GO and KEGG analyses. Subsequently, the candidate genes related to cell proliferation were obtained (DMTF1 and WDR5), which were considered to possibly be related to the slow growth and development of Shenxian pigs. In addition, the candidate genes related to lactation were obtained (CSN2 and CSN3).

## INTRODUCTION

China has a long history and rich experience in livestock and poultry breeding, and it is also one of the main domestication origins of pigs(Giuffra et al.2000;Groenen et al.2012). China has a vast territory, complex terrain, and rich pig resource pool. It can be seen that even the southern China pig breeds are quite different from the northern China ones(Ai et al.2015). As the only local black pig breed of Hebei Province, the Shenxian pig variety was declared extinct in 2004 due to the introduction of a large number of foreign pig breeds. Therefore, in order to protect and study the Shenxian pig breed, Hebei Zhengnong Animal Husbandry Co., Ltd. was established in Hebei Province, with the goal of preserving the purified Shenxian pig variety. Shenxian pigs were listed in the Genetics of Livestock and Poultry Resources of China in 2016. At the present time, a preserved population of Shenxian pigs has been developed, which consists of 28 boars and 220 sows. As a local breed in China, Shenxian pigs are known to have the excellent qualities of the majority of local pig breeds, with relatively strong adaptability to the environmental conditions and resistance to crude feed. The Shenxian pigs have been observed to have stronger reproductive capacities than introduced breeds, with the advantageous characteristics of early sexual maturity and larger litter sizes, and sows with strong lactation abilities. In addition, the breed is known for its higher intramuscular fat content and superior meat quality when compared with other domestic breeds. At the present time, there have been few studies conducted regarding the molecular aspects of the Shenxian pig variety. In recent years, several research studies and tests have been carried out for Shenxian pigs using sequencing methods and other means. For example, using genome-wide association analyses of the main reproductive traits of Shenxian pigs, significant SNP sites which affect the reproductive traits of Shenxian pigs have been successfully screened. In addition, following various post-processing and software analysis processes, some significant SNP sites have been identified which affect the reproductive traits of the breed, and relevant target genes were obtained through site annotation. The previous related studies have identified TSKU, LRRC32, B3GNT6, WNT, LRRC32, CAPN5, MYO7A, TCHHL1, SI00A11, TACR3, IL12A, and THEM5 as the potential candidate genes which affect the reproductive performances of Shenxian pigs. In the present study, based on newly developed resequencing technology, ten Shenxian pigs, nine large white pigs, and ten Meishan pigs were subjected to genetic analyses of population in order to obtain a systematic understanding regarding the population genetic relationships of Shenxian pigs. At the same time, a selection signal study was carried out with the large white pig genome, and selected regions in the genome were compared and analyzed. The goals were to deepen the current understanding of the selection differences between the Shenxian and large white pig breeds, as well as to identify the selection regions related to meat quality and reproductive traits, with the long-term goal of accelerating the breeding processes of the Shenxian pig variety.

## MATERIALS AND METHODS

### Materials

The Shenxian pigs used in this experiment were selected from Hebei Zhengnong Animal Husbandry Co., Ltd., from the resource group of 150 sows from 10 families of Shenxian pigs, 10 21-day-old healthy and newly weaned piglets were selected from each family. Pick one of each and use ear tag tongs to collect the ear tissues of pigs in Shenxian County. Before collection, cut the ear tissues and sterilize them with 75% alcohol. Use ear tag tongs to punch and sample the pig ears. Take about 100mg samples. The 95% alcohol EP tube is transported back to the laboratory in an ice bag and stored in a refrigerator at −20 °C for later use. During sampling, transportation and storage, ensure that the DNA sample is not contaminated and used for subsequent genomic DNA extraction.

### Whole genome resequencing

Whole genome resequencing is process used to sequence the genomes of different individuals of species with known genome sequences, and then analyze the differences of individuals or groups on that basis(Ley et al.2008). It has been found that based on whole genome resequencing technology, researchers have been able to quickly carry out resource surveys and screening processes in order to determine large numbers of genetic variations, and realize genetic evolution analyses and predictions of important candidate genes. In the current study, the Shenxian pigs were re-sequenced for the purpose of obtaining the genomic information. A large number of high accuracy SNPs, InDel, and other variation infor-mation were obtained by comparing the results with the reference genome. Subsequently, the variation information was successfully detected, annotated, and counted.

### Detection methods based on next generation sequencing technology

At present, the second-generation sequencing technology is widely used(Levy et al.2016;Slatko et al.2018). The development of the next generation sequencing (NGS) technology has resulted in revolutionary changes in the detections of genetic variations. Researchers can now comprehensively and accurately detect various types and the sizes of genetic variations from the genomic level(Mardis.2008) For the detection of genetic variations based on the next generation sequencing technology, the first step is to compare the reading segments of thousands of sequences to the corresponding positions on the reference genome. Then, the possible variation information can be inferred using simple mathe-matical models(Andersson.2009;Mckenna et al.2010;Montgomery et al.2013). These methods take the number of reading segments supporting the variations and reference sequences; quality of the reading segment comparison; environmental conditions of the genome sequences; and the possible noise (such as sequencing errors) as prior information. Then, the variation types and genotypes at certain positions can be determined according to naive Bayesian theory, and the genotype with the highest posterior probability can be successfully obtained(Montgomery et al.2013;Durbin et al.2008). Such methods have been found to be effective for SNP and small InDel detections(Depristo et al.2011;Albers et al.2011).

### Data processing

This test sample uses the whole genome resequencing data (sequencing depth 10X) of 10 Shenxian pigs (5 cucumber-mouthed pigs and 5 Wuhuatou pigs). The output of the genome data comparison is the sam file. Since the sorting method of the sam file cannot be used for subsequent analysis, the Samtools software(Li et al.2009)is used to convert the sam file into a file in chromosome sorting format, that is, the bam format. Chromosome rearrangement is to arrange the entries of the same chromosome in the file in ascending order, and merge the two files of the same sample.Download the resequencing genome data of 9 large white pigs and 10 Meishan pigs from the SRA database of the NCBI database (http://www.ncbi.nlm.nih.gov/sra). The downloaded file is generally in sra format and needs to be used The SRA Toolkit tool converts the downloaded sequencing data into a fastq format file for SNP identification.

### SNP detection and annotation

Use GATK and VarScan software(Koboldt et al.2009)to perform SNP detection at the same time, so as to ensure that the obtained SNP site information will not be affected by the deviation of base misalignment caused by InDel mutation. GATK detection SNP code: java -jar GenomeAnalysisTK.jar glm SNP -R ref.fa -T UnifiedGenotyper -I test.sorted.repeatmark.bam -o test.raw.vcf. In order to ensure the accuracy of SNP information, strict testing conditions are set when detecting SNP site information: <1>The minimum number of end reads greater or equal to 4; <2>The minimum quality value Q20 greater or equal to 90; <3>Minimum coverage greater or equal to 6; <4>P value Less than or equal to 0.01. InDel detection and annotation: The same use of GATK and VarScan software for InDel detection can eliminate errors caused by SNP variation. GATK detection InDel code: java -jar GenomeAnalysisTK.jar glm InDel -R ref.fa -T UnifiedGenotyper -I test.sorted.repeatmark.bam -o test.raw.vcf. The detection conditions are consistent with SNP. The detected SNP and InDel mutation sites also need quality control, remove untrusted data sites, correct the quality value, and output the file in VCF format after correction. Use R language(Gao et al.2014;Sudhaka.2018)and ANNOVAR software (Kai et al.2014)to perform mutation information Sort and comment.

### Population structure analysis based on SNP

A variety of methods were used to analyze the genetic relationships among the three populations.Plink (V1.90) software was used for principal component analysis (PCA) of shenxian pig, Large white pig and Meishan pig populations, and the top two characteristic vector values were selected for analysis. Generally, the closer the spatial linear distance in the PCA graph was, the closer the genetic relationship of individuals was.Plink (V1.9)(Purcell et al.2007)was used to construct the IBS genetic distance matrix, and phylogenetic trees were constructed by neighbor-joining (NJ) through the genetic distance matrix, and the genetic trees were drawn by mega software.PopLDdecay software was used to analyze the chain imbalance, analyze the correlation of alleles of the three populations, and determine the genetic richness of the population according to the decay rate of LD.Analysis of population genetic structure using Admixture software helps understand evolutionary processes, differentiate migrating individuals and hybrids, etc., and identify subgroups to which individuals belong through genotype and phenotypic association studies.

## RESULTS AND DISCUSSION

### Reference genome alignment

BWA software (Version: 0.7.15-r1140) was used in this research investigation to compare the clean sequences of the two subpopulations of Shenxian pigs after quality control with the latest version of reference genome of pigs was implemented. The results are detailed in Table 1. It was found that on average, more than 86 percent of the sequencing data could be compared to the reference genome.

**Table 1.** Reference genome comparison results statistics

| Sample-name<sup>a</sup> | Mapped<sup>b</sup> | Paired<sup>c</sup> | Insert<sup>d</sup> | SD<sup>e</sup> |
| --- | --- | --- | --- | --- |
| B1-1 | 86.75 | 76.36 | 387 | '-128/+129 |
| B1-2 | 86.74 | 74.70 | 408 | '-129/+134 |
| B1-3 | 87.12 | 75.61 | 407 | '-124/+124 |
| B1-4 | 87.09 | 76.07 | 415 | '-126/+124 |
| B1-5 | 87.01 | 75.91 | 365 | '-110/+114 |
| B2-1 | 86.11 | 75.40 | 399 | '-120/+115 |
| B2-2 | 85.02 | 73.37 | 396 | '-122/+122 |
| B2-3 | 87.15 | 75.39 | 386 | '-118/+135 |
| B2-4 | 87.04 | 75.15 | 436 | '-141/+140 |
| B2-5 | 86.66 | 75.44 | 380 | '-114/+120 |

<sup>a</sup> Huangguazui:B1,Wuhuatou:B2 <sup>b</sup> The proportion of valid data that can be compared to the reads on the reference genome <sup>c</sup> Both read1 and read2 can compare the ratio of reads on the reference genome to valid data <sup>d</sup> The length of the insert, the unit is bp <sup>e</sup> Compare the mean square error of reads

### SNP identification

Clean data were obtained after strict quality control and alignment processes of the obtained genome data were implemented. Then, GATK software was used to identify the SNP, and the SNP data were used for the subsequent analysis processes. A total of 34.2M SNP sites were identified in the examined Shenxian, large white, and Meishan pig individuals. There were determined to be 23M SNP sites in the Shenxian pig and 8.8M SNP sites in the large white pig varieties. In addition, there were 2.2M SNP sites in Meishan pig, and a total of 6.9M SNP sites in Shenxian pig and large white pig, respectively. This study found that more SNP sites had been identified in Shenxian pig strain, while a total of 1.5M SNP sites were identified in the Shenxian, large white, and Meishan pig varieties. It was possible that Shenxian pig breed had shown more genetic variations in the pig reference genome, while the Meishan and large white pig varieties had displayed less variability with the identified SNP sites when compared with the Shenxian pig breed (Table 2). However, it was found to be difficult to extract common sites for analysis purposes, and the Meishan pigs were observed to be missing sites during the extraction processes of the common sites. Therefore, the subsequent population structure analyses and LD attenuation analyses were analyzed in pairs.

**Table 2.**
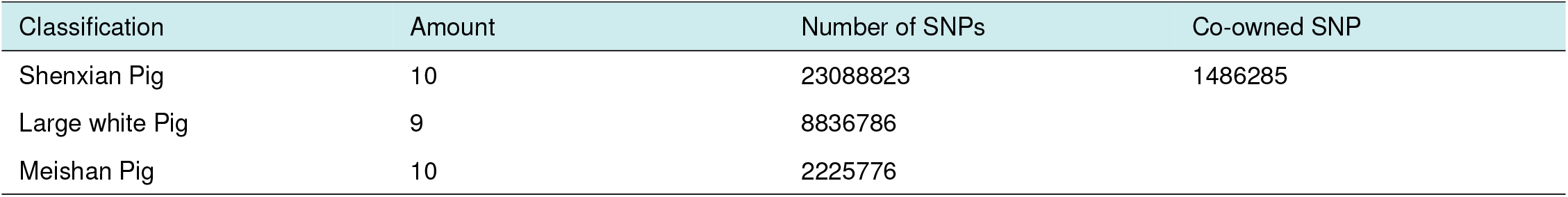
Statistics of the SNP results

## POPULATION STRUCTURE ANALYSES

### Principal component analyses

In order to reveal the variety specificity of Shenxian pigs, principal component analyses (PCA) were performed on the obtained SNP information of the Shenxian, large white, and Meishan pig breeds, respectively, as shown in Figure 1 to Figure 3). Then, the first three eigenvectors were selected. The analysis results of the first eigenvector and the second eigenvector revealed that the first eigenvector had effectively distinguished the three breeds. The second eigenvector showed that the large white pig population was a population with small differentiations within the population. Meanwhile, population stratification was observed within the Shenxian and Meishan pig populations. The analysis results of the first eigenvector and the third eigenvector showed that the population stratification of the large white pig and Shenxian pig breeds was not obvious. However, the Meishan pig population displayed obvious stratification. The second eigenvector and the third eigenvector also revealed that there was a certain degree of stratification in the Meishan pig population. Furthermore, the results indicated that the stratification in the Meishan pig population was larger than that in both the Shenxian pig and large white pig populations.

**Figure 1.**
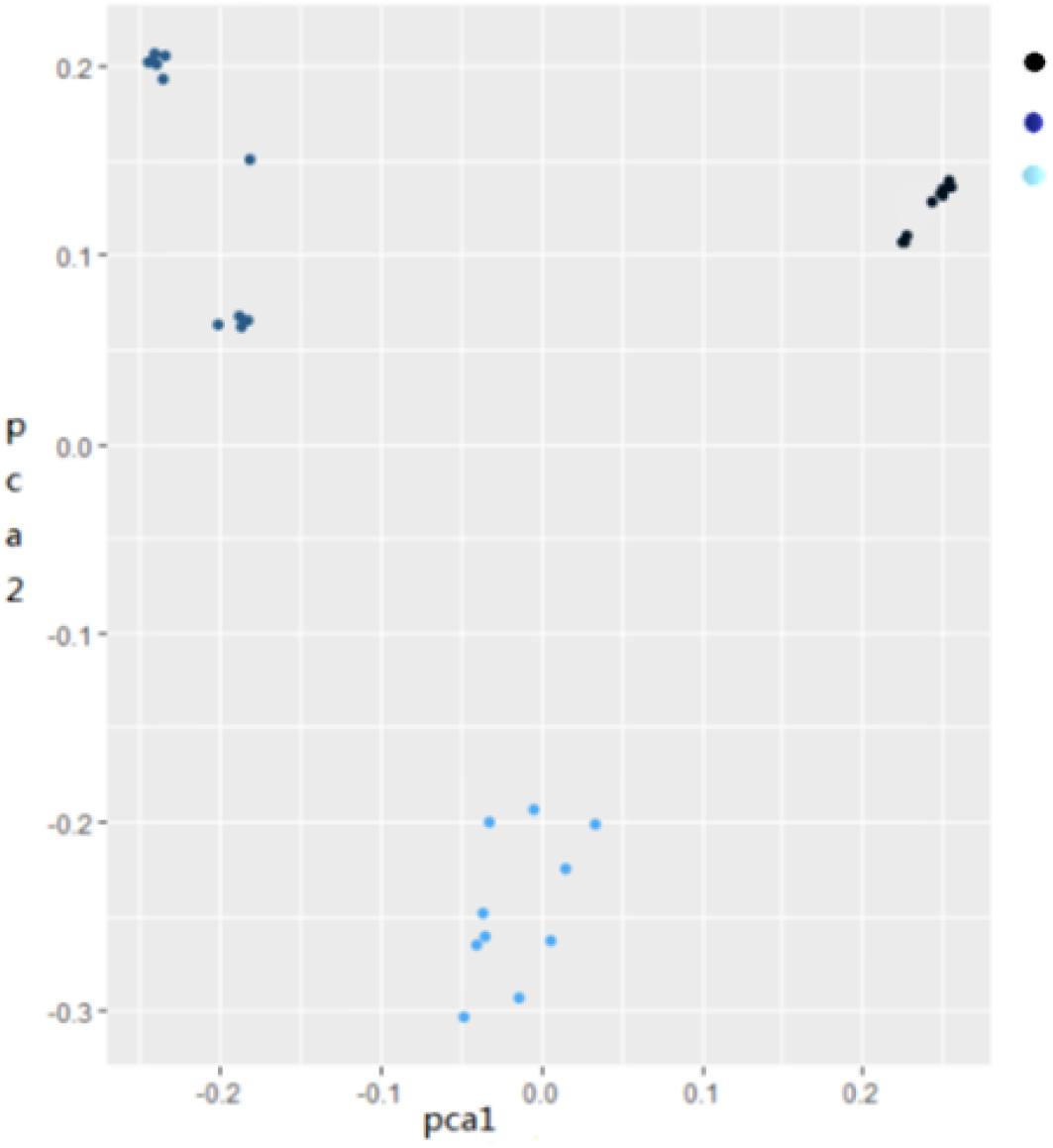
The analysis of the first feature vector and the second feature vector shows that the first vector vector distinguishes the three breeds well, and the second feature vector shows that the large white pig population is one breed.

**Figure 2.**
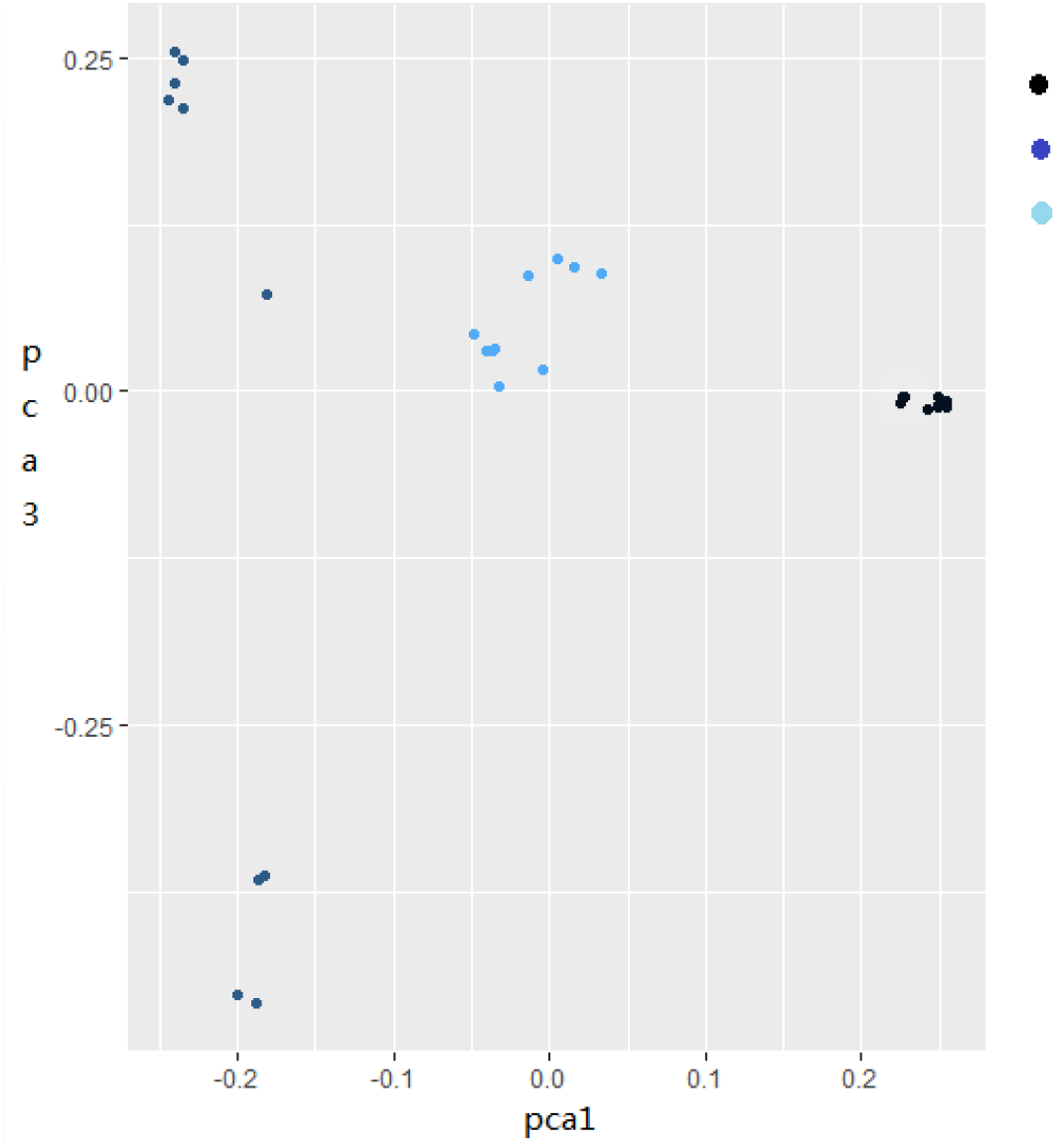
The analysis of the first eigenvector and the third eigenvector shows that the large white pig and Shenxian pig populations are not clearly stratified, while the Meishan pig population has obvious stratification.

**Figure 3.**
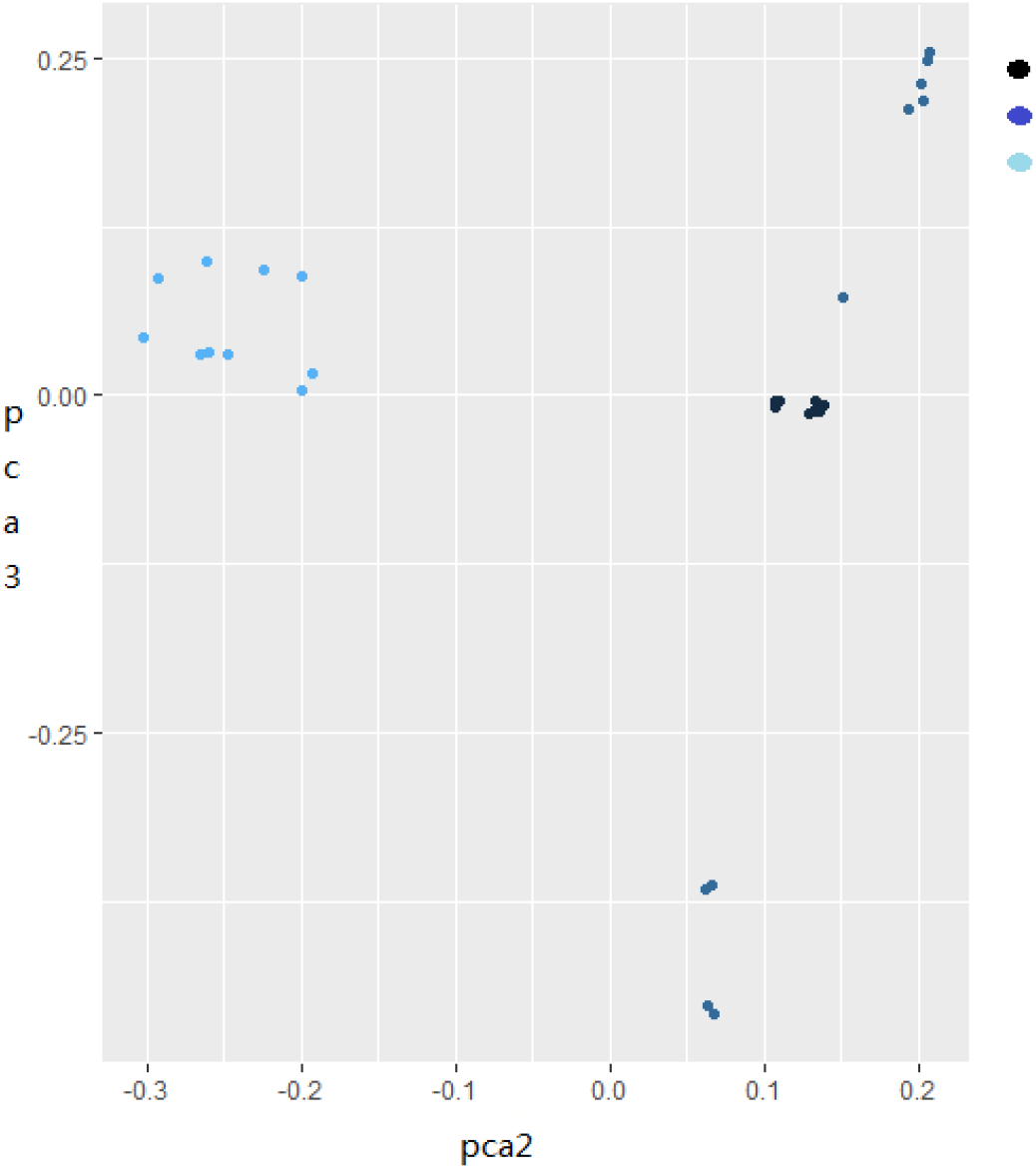
The analysis of the second feature vector and the third feature vector shows that the Meishan pig population has a certain degree of stratification.

### Phylogenetic tree analyses

In order to verify the results of the principal component analyses (PCA) among the Shenxian, large white, and Meishan pig breeds, the SNP data were used to construct the genetic distance matrixes of the three populations in plink software. Then, an evolutionary relationship analysis was carried out in order to draw the evolutionary tree, as shown in Figure 4. The analysis results showed that the three populations had originated from the same species. In addition, there were distant genetic relationships observed between the Shenxian pigs and the other two populations. However, when compared with the Shenxian pigs, the Meishan pigs were found to have a closer genetic relationship with the large white pigs.

**Figure 4.**
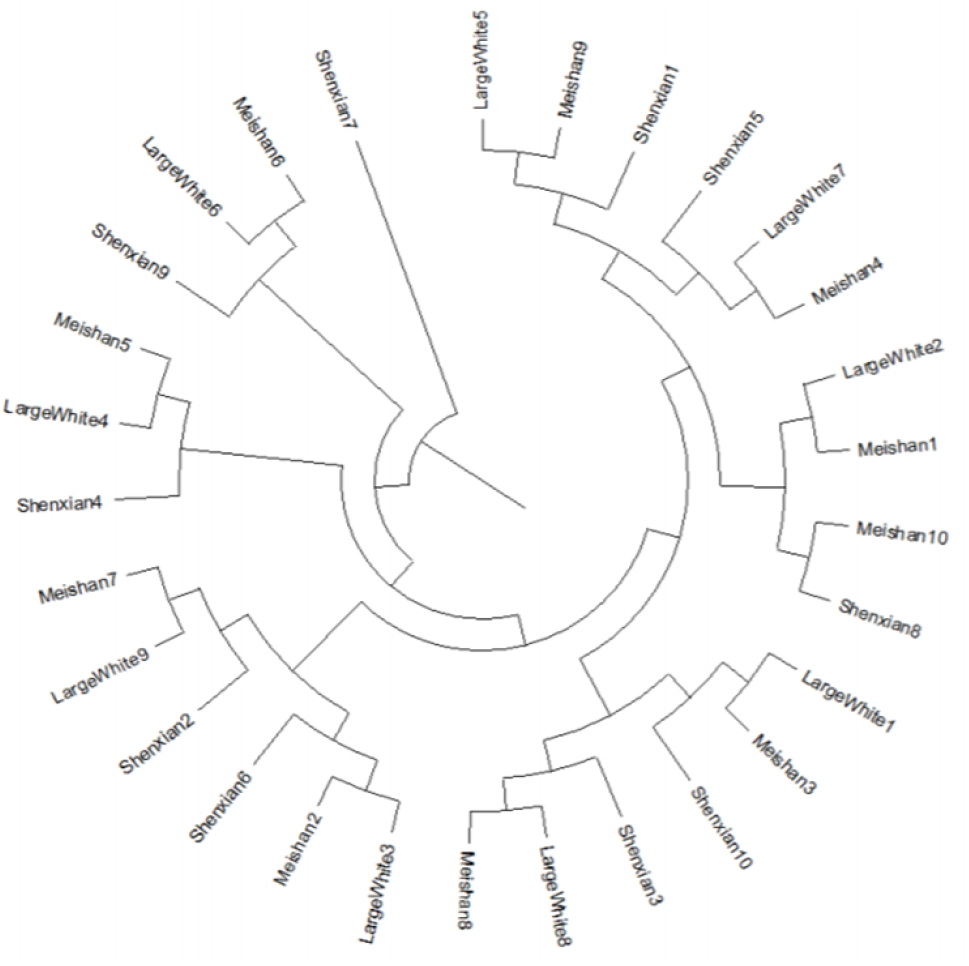
The NJtree showed that the three populations had originated from the same species. In addition, there were distant genetic relationships observed between the Shenxian pigs and the other two populations. However, when compared with the Shenxian pigs, the Meishan pigs were found to have a closer genetic relationship with the large white pigs.

### Population structure analyses

Previous studies have shown that Asian breeds of pigs had completed domestication earlier than European breeds. However, due to geographic migrations, the genomes of European domesticated pigs are likely to include a large number of Asian pig genomes. In the current study, in order to explore the ancestral relationships of Shenxian pigs, the population structures of the Shenxian pig, large white varieties were further analyzed, as detailed in Figure 5. It was observed that the degrees of intergenomic communication between the Shenxian pigs and the large white pigs were basically consistent. there were observed to be relatively more gene exchanges between the Shenxian and the large white pigs. These difference may have been caused by many factors, such as the infiltration of the Shenxian pig genes into European domesticated pigs after domestication was complete, or the introduction of the genome of the large white pig during the development and breed conservation of the Shenxian pig breed. Another possibility may have been the existence of common genomes between the two breeds.

**Figure 5.**
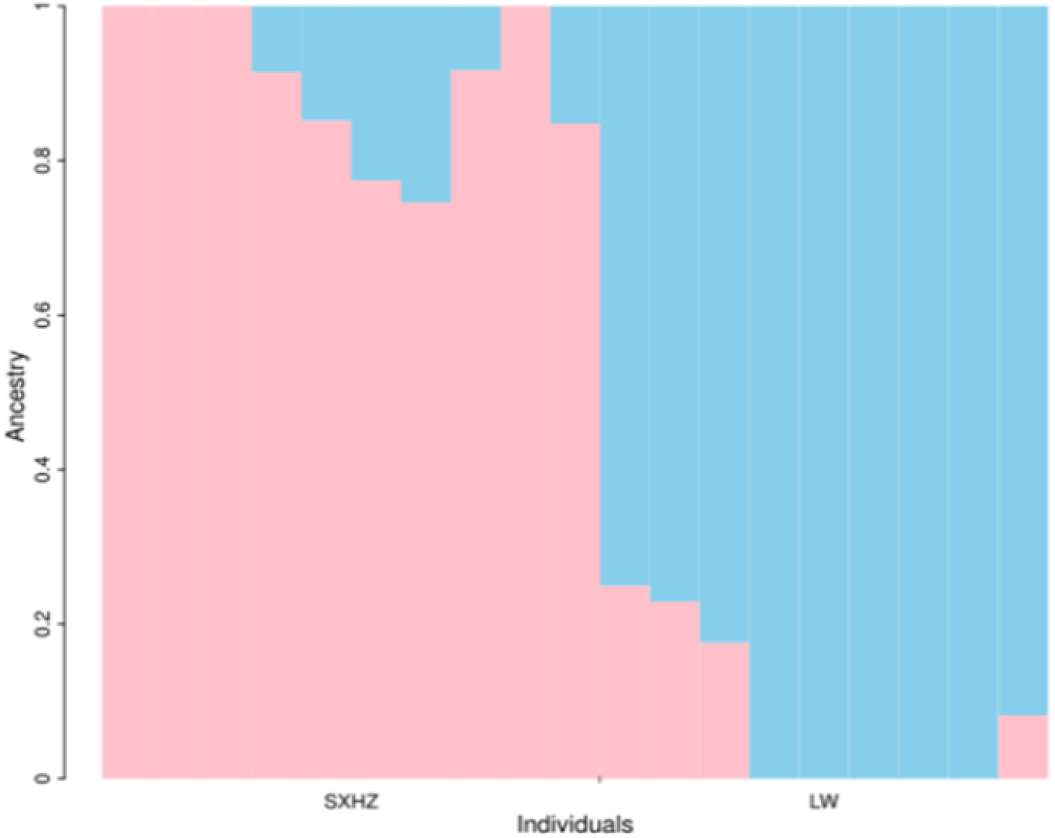
There are relatively more gene exchanges between Shenxian pigs and Dabai. Shenxian pigs may have introduced Dabai’s genome in the development and conservation of breeds, or they may share a common genome.

**Figure 6.**
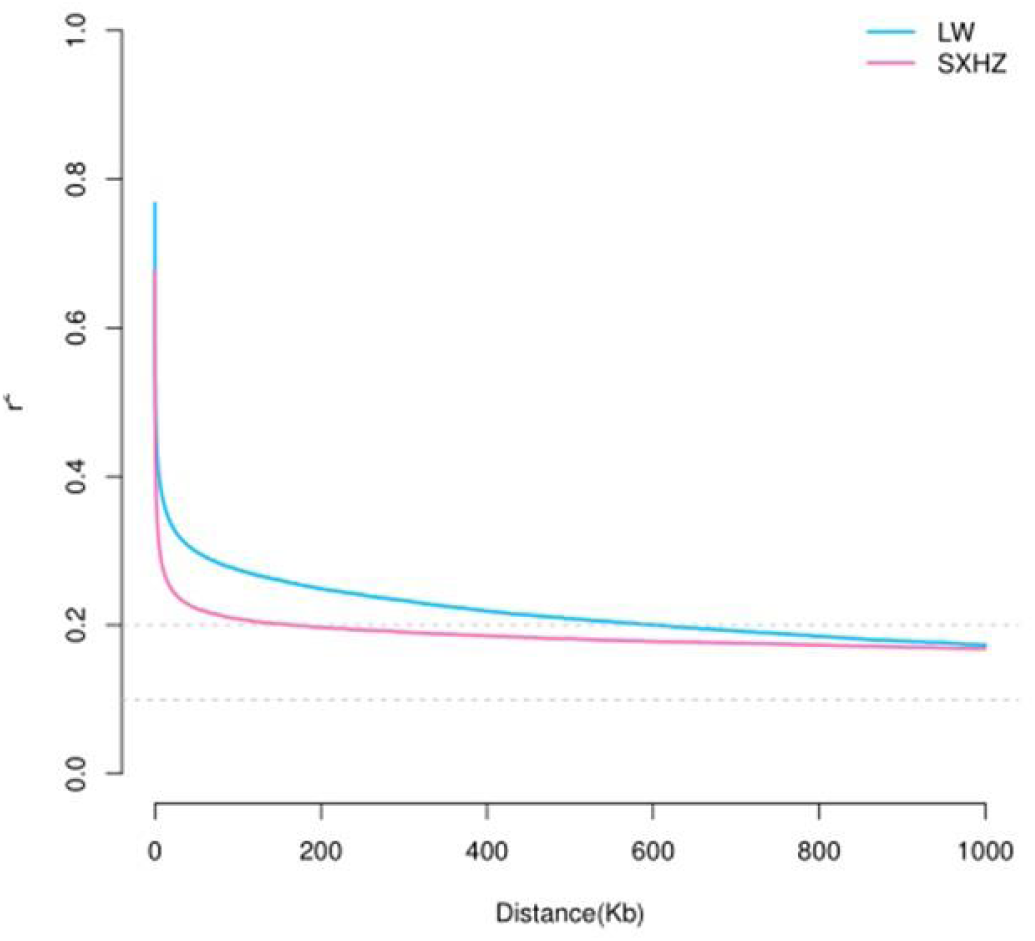
Generally speaking, the higher the domestication intensity was, the greater the selection intensity would be, and the smaller the LD attenuation rate would be. The results shown in the LD attenuation diagram (Figure 6) revealed that the LD coefficient of the Shenxian pig population declined faster than that of the large white pig population. These findings indicated that the genetic diversity of the Shenxian pig population was higher.

### Analysis results of the LD (linkage disequilibrium) attenuation

The attenuation rates of the LD were found to be different among the different pig populations. It was found that the higher the selection intensity was, the slower the attenuation rate of the LD would be. The linkage disequilibrium of the Shenxian, large white populations was analyzed using PopLDdecay software. It was determined from the LD attenuation map that the linkage disequilibrium of the different populations decreased with the increases in marker spacings. The value of D was found to be strongly dependent on the artificial allele frequency, which was not conducive to a comparison of the LD attenuation. Therefore, the standardized disequilibrium coefficient D, was adopted in order to avoid the dependence on the allele frequency. The calculation method of D’ was as follows: D’ = D/Dmax When D<0,Dmax = minP(A)P(B), P(a)P(b); When D<0,Dmax = minP(A)P(b), P(a)P(B); When D’ = 1, it was indicated that the linkage was totally unbalanced and there was no reorganization; Finally, when D’ = 0, it was indicated that the linkage was completely balanced with random reorganization.

### Fst and Nucleic acid diversity analyses

In order to study the selection preferences of the Shenxian pigs, this study analyzed the selection signals of both the large white pigs and the Shenxian pigs. Ten Shenxian pigs forming a population, and nine large white pigs forming a population, were selected for this study’s signal analysis process. The detection strategy of combining population differentiation index Fst and nucleic acid diversity Nucleic acid diversity was adopted. A population differentiation index (Fst) method was used to anchor the regions of the genome differentiation between the populations. A nucleic acid diversity method was used to anchor the regions of the reduced polymorphism within the populations. The regions identified by the two methods were selected as the candidate regions for the purpose of improving the reliability of the results. In addition, in order to reduce the impacts of any outliers, this study utilized a sliding window detection method. The sliding window size was set as 10 kbp and the sliding window step was set as 1 kbp. In order to compare the results of the two analyses, the Fst values and the Nucleic acid diversity ratio (Nucleic acid diversity large white pig/ Nucleic acid diversity Shenxian pig) values were adjusted to a unified scale using a minimum/maximum conversion process, as detailed in Figure 7.

**Figure 7.**
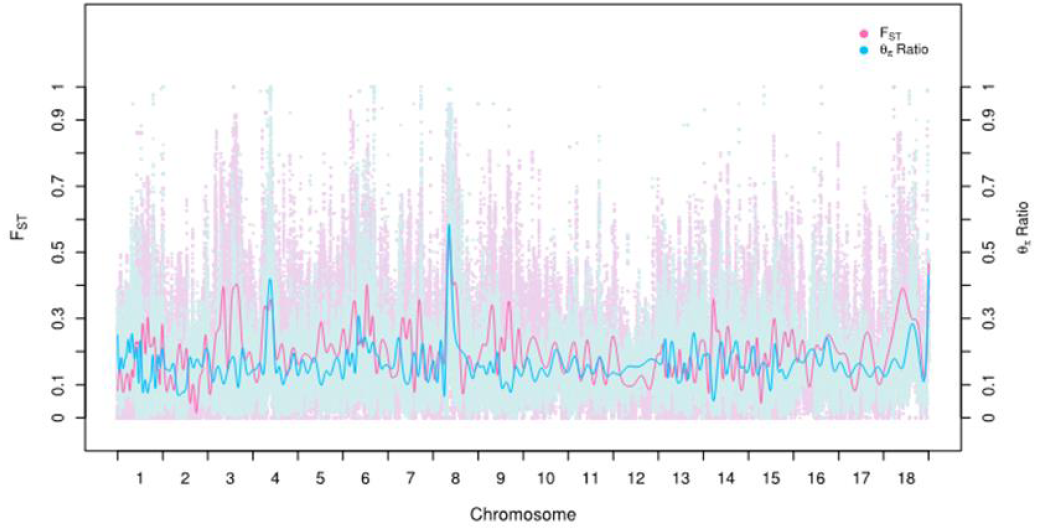
Figure 7 shows the results of Fst and Nucleic acid diversity following the completions of size conversions; Red represents the results of the Fst and blue represents the results of Nucleic acid diversity ratio

### Inference of the selected sites

The top 1% window of the Fst slide window analysis and the top 1% window of the Nucleic acid diversity analysis were screened in order to obtain 1,509 selected sites, and the selected sites were then annotated. Figure 8 and Figure 9, respectively, show the distributions of the selected variation sites in the chromosome and genome. Figure 8 shows that when compared with large white pigs, the Shenxian pigs had displayed more selection variations on chromosome 8 and less variations on the other chromosomes. As can be seen from Figure 9, the selection of those sites was mainly concentrated between the intergenic region and the intron, while the selection in other regions was less. However, the variations of the intergenic regions and intron had little effect on the gene expressions, which may be due to the fact that the changes in selection pressure after species differentiation had resulted in major differences between intron and intergenic regions.

**Figure 8.**
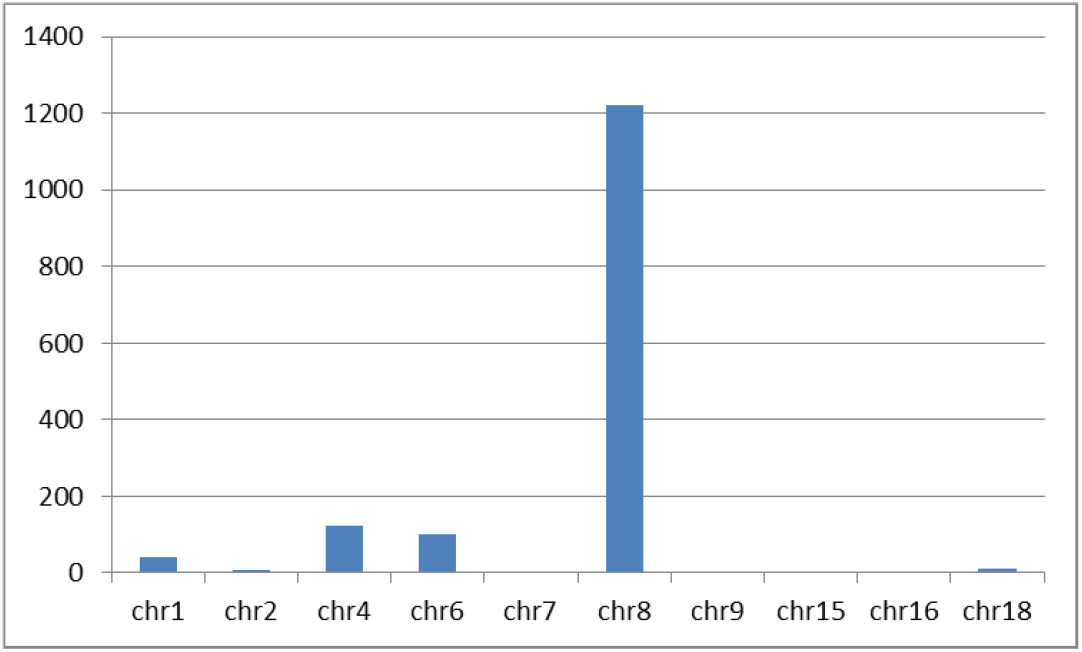
Figure 8 Distribution of the variations on the chromosomes. Note: In the figure, the abscissa represents the chromosome number; and the ordinate represents the number of selected sites.

**Figure 9.**
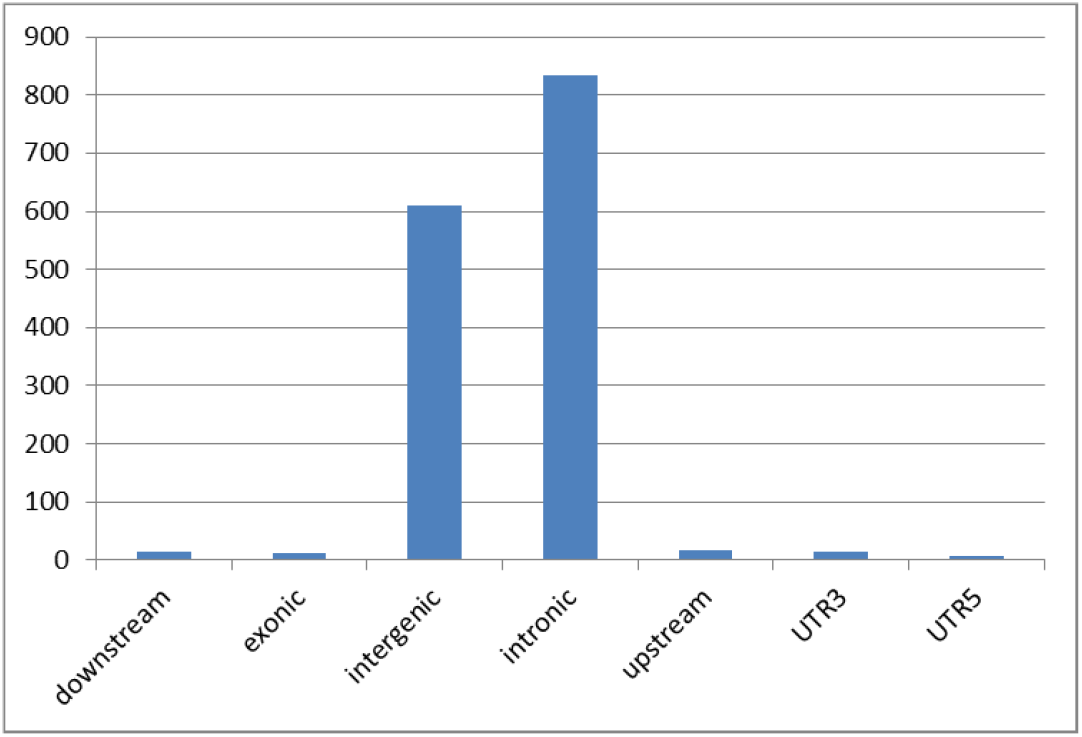
Figure 9 Distribution of the variation locations.Note: In the figure, the abscissa represents the gene region; and the ordinate represents the number of selected sites

### Functional enrichment analysis of the candidate gene GO

In the current research investigation, the 19 differentially expressed genes at 1,509 sites obtained through the screening process were subjected to functional enrichment and significance testing processes. The GO enrichment consisted of a total of three parts(Figure 10 to Figure 12): biological process (BP);cell composition (CC) and Molecular function (MF), which were enriched to 421 entries. Then, through the enrichment analyses of the genes, the biological processes involved in those genes were successfully identified. From the result, the selection regions of the Shenxian pigs relative to the large white pigs during the processes of genetic evolution and domestication could be identified, and the selection characteristics of the Shenxian pig population were more clearly understood.

**Figure 10.**
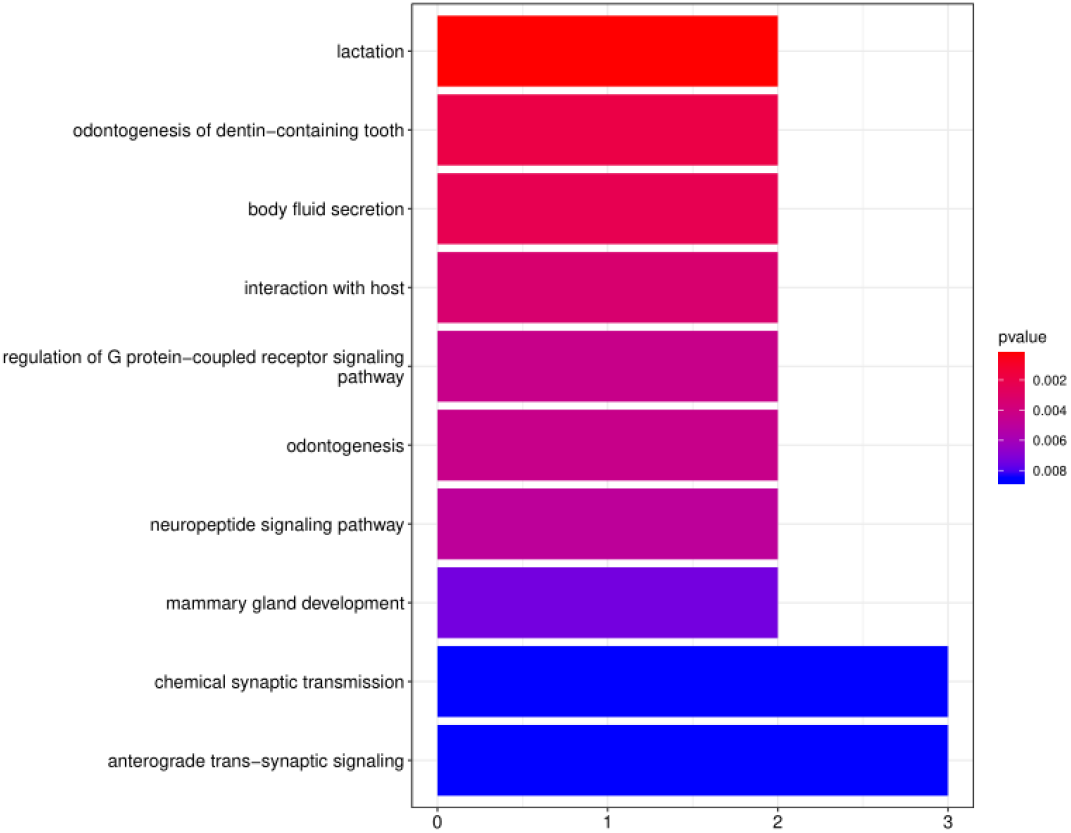
The main involved biological processes included lactation; body fluid secretion; host interaction; regulatory pathways of the G protein coupled receptor signal transduction; dentinogenesis; neuropeptide signaling pathway; breast development; chemical synaptic transmission; antegrade synaptic signaling, and so on.

**Figure 11.**
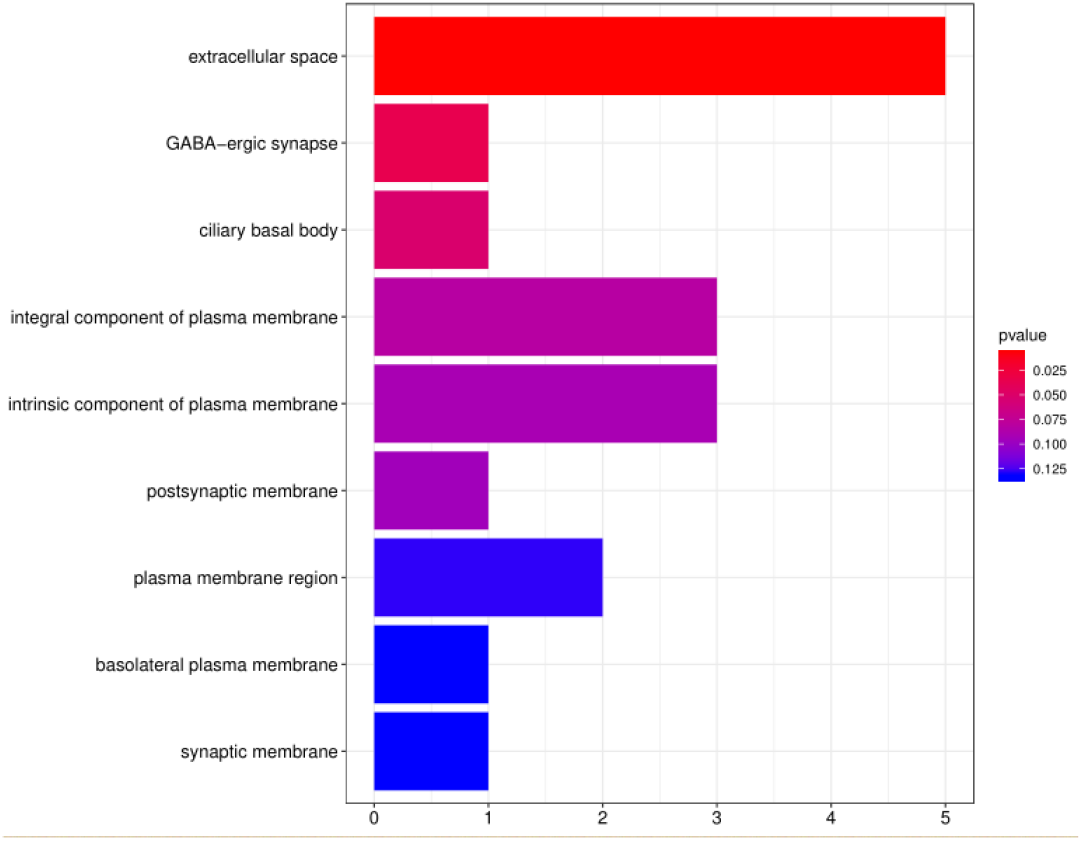
The cell compositions indicated that the candidate genes may also participate in extracellular space; synapse; ciliated body; protoplasm membrane composition; plasma membrane composition; postsynaptic membrane; plasma membrane interval; basal outer plasma membrane; and synaptic membrane processes.

**Figure 12.**
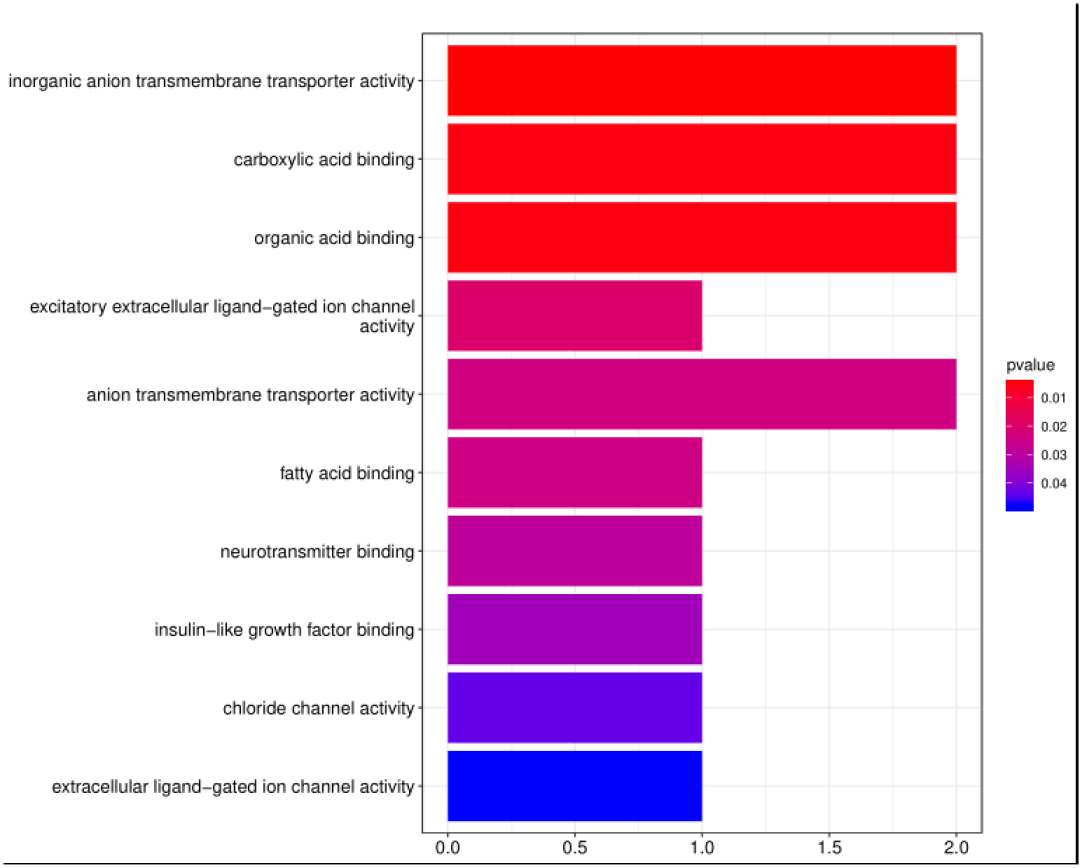
The molecular function enrichment results revealed that the candidate genes were involved in inorganic anion transmembrane transporter activities; anion transmembrane transporter activities; hydroxyl acid binding; organic acid binding; fatty acid binding; excitatory cell ligand ion channel activities; neurotransmitter binding; insulin-like growth factor binding; chloride ion binding; cell apoptosis Ligand ion channel activities, and other molecular functions.

**Figure 13.**
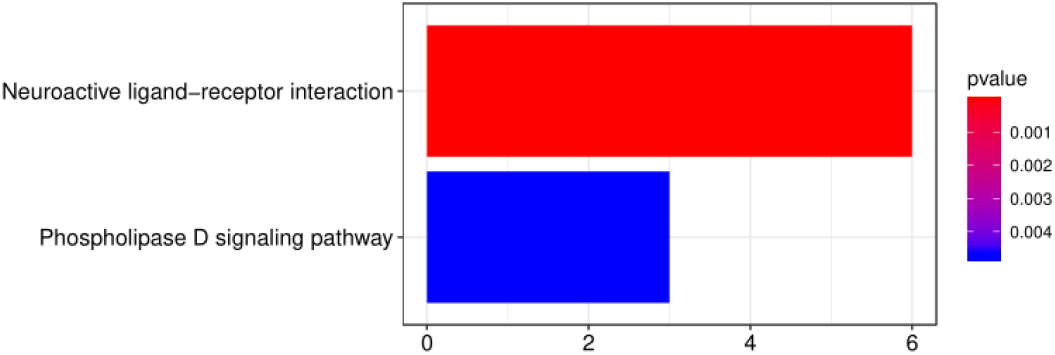
In the figure, the enriched KEGG pathway terms include the horizontal axis representing the number of genes, and the vertical axis representing the enriched pathways

### Genes significantly enriched into pathways

Through the KEGG pathway analyses of the candidate genes revealed that those genes were mainly involved in two pathways, as detailed in Figure 3. It was found that among the 19 genes, six genes were involved in the neuroactive receptor interaction pathways, and three genes were involved in phospholipase D signaling pathways(Table 3).

**Table 3.** Detailed list of pathway enriched genes

| ID | Description | GeneRatio | Pvalue | GeneID |
| --- | --- | --- | --- | --- |
| ssc04080 | Neuroactiveligand-receptor interaction | 6/19 | 8.91142E-05 | 100152093/414284/100622950<br>100623130/397547/100524120 |
| ssc04072 | PhospholipaseD signaling pathway | 3/19 | 0.004736493 | 100515267/396880/100524120 |

This experiment analyzed the selection areas of the local breed Shenxian pig and the Western commercial breed Large White pig. The results showed that the two breeds have the same artificial selection in some signaling pathways, but most of the selection areas are different, natural selection and artificial selection. The difference in selection may have resulted in two different breeds. Large white pigs have developed into more mature commercial breeds and have been promoted all over the world. Shenxian pigs, as a developing local breed, are designed to preserve the individual characteristics and characteristics of Shenxian pigs. The selection traces of excellent genes need further study. The study only conducted the research on the breed selection signal of the Shenxian pig and the Large White pig, and learned about the selection characteristics of the introduced breed of the Shenxian pig, but it did not have a good understanding of the genetic characteristics of the Shenxian pig as a local breed in China. Complete selection signal analysis of Shenxian pigs and different breeds, analysis of selection signals with wild populations to understand the traces of artificial selection of Shenxian pigs, and selection signal analysis of other local breeds to understand the selection characteristics of different breeds and dig out functional genes In order to truly understand the breed diversity and group characteristics of pigs in Shenxian County.

## Discussion

In this study, the screened genes included CXCL8, GLRB, CD180, TMOD2, AFP, ALB, RASSF6, AMBN, CSN2, CSN3, DMTF1, IGFBP7, IMPAD1, NAP1l4, NPY1R, ODAM, SLC4A4, WDR5, and AFM. Among those genes, CXCL8(Wang et al.2020;UkaszewiczZajc et al.2020), CD180(Dong et al.2019), AFP(B et al.2020), RASSF6(Vogel et al.2020), and IGFBP7 have been proved to be related to immune diseases, and these genes are closely related to human diseases. For example, CXCL8 is a chemokine which is expressed in a variety of tissues. In human studies, it has been found that the CXCL8 gene is related to cancer, and its expression levels in cancer tissue are higher than those in other tissue(Tongbo et al.2019). At the same time, it may also cause hypertension (Young et al.2018), and its expression levels have been found to be higher in patients with chronic hepatitis B (Sharif et al.2018). Furthermore, this gene can cause inflammatory reactions in pigs(Hu et al.2011). It is of major significance to study this gene for disease research purposes. In previous studies of diseases, CD180 was found to be related to systemic lupus erythematosus. The expression levels of CD180 in the cells of patients with systemic lupus erythematosus were higher than those of patients without systemic lupus erythematosus. At the same time, CD180 is considered to be related to chronic lymphoproliferative diseases(Jing et al.2018). However, this gene has not been reported in previous studies regarding pigs. The AFP gene is related to human liver cancer, lung cancer, pancreas, and other diseases, and the research regarding AFP in pigs is also rare(Kim et al.2003). The RASSF6 gene is known to be associated with colon cancer disease, and can promote the growth of colon cancer cells(Wang et al.2020). The IGFBP7 gene(Zhang et al.2019;Chenyi et al.2020) has been determined to be related to osteoarthritis. The gene expression levels in cartilage tissue of patients with osteoarthritis have been found to be high, and the gene can accelerate chondrocyte apoptosis by inhibiting chondrocyte proliferation. The DMTF1(Tschan et al.2015;Yang et al.2018), WDR5, and other genes and known to be related to cell proliferation. The DMTF1 gene is also related to cancer diseases(Peng et al.2015). The WDR5 gene plays a certain role in tissue regeneration and bone tissue development(B et al.2020), and this gene and family protein have been confirmed to be able to maintain the normal development of pig preimplantation embryos(Gallenberger et al.2011;Maserati et al.2011;Ye et al.2011). The CSN2(Suteu et al.2019;Kumar et al.2019)and CSN3(Curi et al.2005)genes have been found to be related to lactation. At the same time, there were also other genes observed in the selected regions. For example, the AMBN gene may be related to tooth development(Ting et al.2018); NAP1l4 gene may be related to cell regulation(Tanaka et al.2019); and the IMPAD1 gene may be related to bone development(Vissers et al.2011). However, there have been few studies conducted regarding the aforementioned genes in pig breeds.

